# On the correspondence between the transcriptomic response of a compound and its effects on its targets

**DOI:** 10.1101/2023.01.01.522411

**Authors:** Chloe Engler Hart, Daniel Ence, David Healey, Daniel Domingo-Fernández

## Abstract

Better understanding the transcriptomic response produced by a compound perturbing its targets can shed light on the underlying biological processes regulated by the compound. However, establishing the relationship between the induced transcriptomic response and the target of a compound is non-trivial, partly because targets are rarely differentially expressed. Thus, connecting both modalities requires orthogonal information (e.g., pathway or functional information). Here, we present a comprehensive study aimed at exploring this relationship by leveraging thousands of transcriptomic experiments and target data for over 2,000 compounds. Firstly, we confirmed that compound-target information does not correlate as expected with the transcriptomic signatures induced by a compound. However, we demonstrate how the concordance between both modalities can be increased by connecting pathway and target information. Additionally, we investigated whether compounds that target the same proteins induce a similar transcriptomic response and conversely, whether compounds with similar transcriptomic responses share the same target proteins. While our findings suggest that this is generally not the case, we did observe that compounds with similar transcriptomic profiles are more likely to share at least one protein target, as well as common therapeutic applications. Lastly, we present a case scenario on a few compound pairs with high similarity to demonstrate how the relationship between both modalities can be exploited for mechanism of action deconvolution.

## 1. Introduction

Transcriptomic data informs about the changes in transcriptional activity in terms of differential mRNA abundance, providing a ‘snapshot’ of cellular signaling. In the last decade, numerous approaches have demonstrated how this information can be leveraged to identify candidate drugs for a given indication (Iorio *et al*., 2010; Sirota *et al*., 2011). Parallely, a vast abundance of bioactivity data has been generated by novel high-throughput techniques that provide information about whether a specific protein is targeted by a chemical compound. Both modalities are complementary, as the binding of the compound to its target(s) leads to transcriptomic changes in genes regulated or modulated by the target (Trapotsi *et al*., 2022).

In recent years, several studies have examined whether transcriptomic signatures can be used to predict the target of a compound. The first study by Isik *et al*. (2015), analyzed over 500 compounds from the Connectivity Map (Lamb *et al*., 2006) and determined that 97% of them did not exhibit any expression changes on their targets in their corresponding drug perturbation experiments. However, they found that dysregulated genes are significantly closer to the compound’s target than by chance by calculating the shortest path between each dysregulated gene and target in a protein-protein interactome. A more recent study by Pabon *et al*. (2018) analyzed the correlation between profiles from drug perturbation and gene knockdown experiments with the L1000 platform. In their work, they evaluated whether looking at the top correlated drug-gene knockdown profiles could identify pairs corresponding to a compound and its target. To do so, they employed 29 compounds for which their target was known, and identified eight true positives among the top 100 predicted drug-gene knockdown pairs. Additionally, they found that the number of true positives slightly increased to 10/100 by investigating correlations of the interacting proteins of the target. Similar to the results of Isik *et al*., this larger scale study found that some of the compounds exhibited a low correlation with the knockdown profile of its target.

While these studies have shown that transcriptomic data alone is insufficient to predict the target of a given chemical, they also revealed that leveraging prior knowledge represented as protein-protein interactions can be used to better understand the relation between the transcriptomic signatures of a compound and its known targets. Furthermore, apart from target prediction, better understanding the relationship between these two modalities can help us understand the Mechanism of Action (MoA) of drugs, since compounds that induce a similar transcriptomic signature might share the same target(s) (Rees *et al*., 2016). Lastly, it is currently unclear to what degree structurally similar compounds that target the same proteins also induce similar gene expression profiles.

Recently, a new database called ChemPert integrated protein target information as well as thousands of transcriptomic experiments from over one hundred non-cancer cell types (Zheng *et al*., 2022). Leveraging this resource, we investigated the correspondence between transcriptomic and target data in over 2,000 compounds. To do so, we represented transcriptomic and target data using different approaches and subsequently evaluated their similarity using various correlation metrics. Our results showed that, in line with previous work, targets are rarely differentially expressed in transcriptomic experiments. However, by combining the target information of the compound with pathway data its correlation with its induced transcriptomic response can increase. Furthermore, we found that compounds targeting the same protein do not necessarily induce a similar transcriptomic response in the same cell line. Inversely, we found that compounds with highly similar transcriptomic profiles are more likely to share at least one protein target as well as therapeutic indications. Finally, we present a case scenario where we explored two natural products that exhibited the highest correlation between their transcriptomic and target vectors to demonstrate how this information can be exploited for MoA deconvolution.

## 2. Methodology

**Figure 1.** illustrates the different analyses conducted. All analyses are based on transcriptomic and target data for over two thousand compounds retrieved from ChemPert (Zheng *et al*., 2022) (see subsection 2.1). These two modalities are represented as vectors as we intend to evaluate their correspondence for each compound. In subsection 2.2, we describe the three approaches used to represent each compound. The correlation metrics used to evaluate the agreement between both modalities are outlined in subsection 2.3. The following section (2.4) describes the different correlation analyses outlined in **Figure 1**. Finally, subsection 2.5 outlines the implementation details.

**Figure 1.**
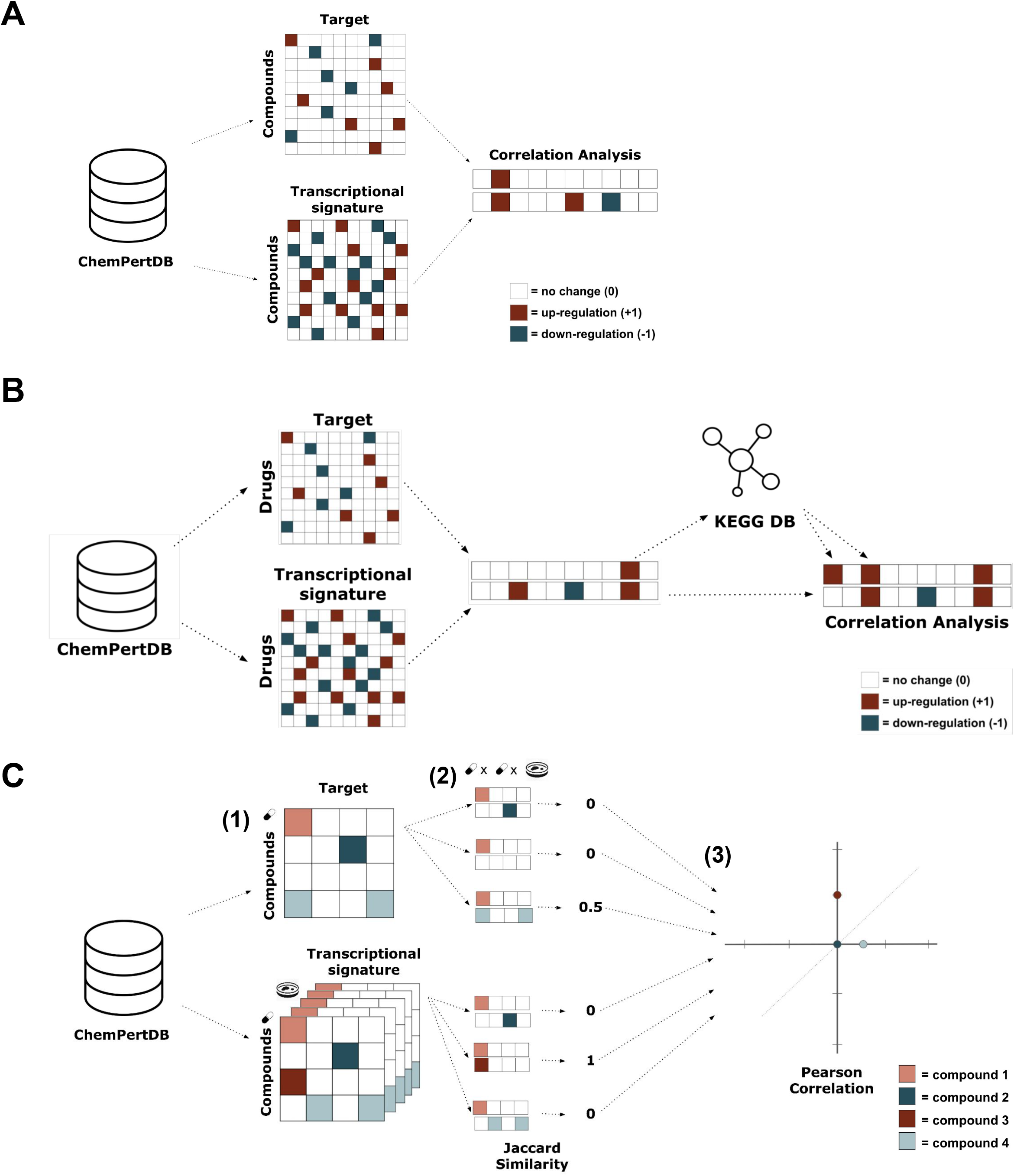

### 2.1. Collecting transcriptomic and target information

We leveraged data from ChemPert (Zheng *et al*., 2022), a database containing transcriptional data from thousands of experiments where cell lines and tissues were perturbed by 2,508 unique perturbagens (details in Supplementary Text 1). Furthermore, this database contains target information from Drug Repurposing Hub (Corsello *et al*., 2017), DrugBank (Wishart *et al*., 2018), and STITCH (Szklarczyk *et al*., 2016). The 82,270 transcriptional signatures at different concentrations of the 2,508 compounds in ChemPert were filtered to a subset of 2,152 chemical compounds for which both transcriptomic and target data was available on 2022-05-04 (**Figure 2D)**. This subset is majorly composed of small molecules and a few peptides. Distributions of the different concentrations measured for each gene, number of Differentially Expressed Genes (DEG), and number of targets for each chemical are shown in **Figure 2**.

**Figure 2.**
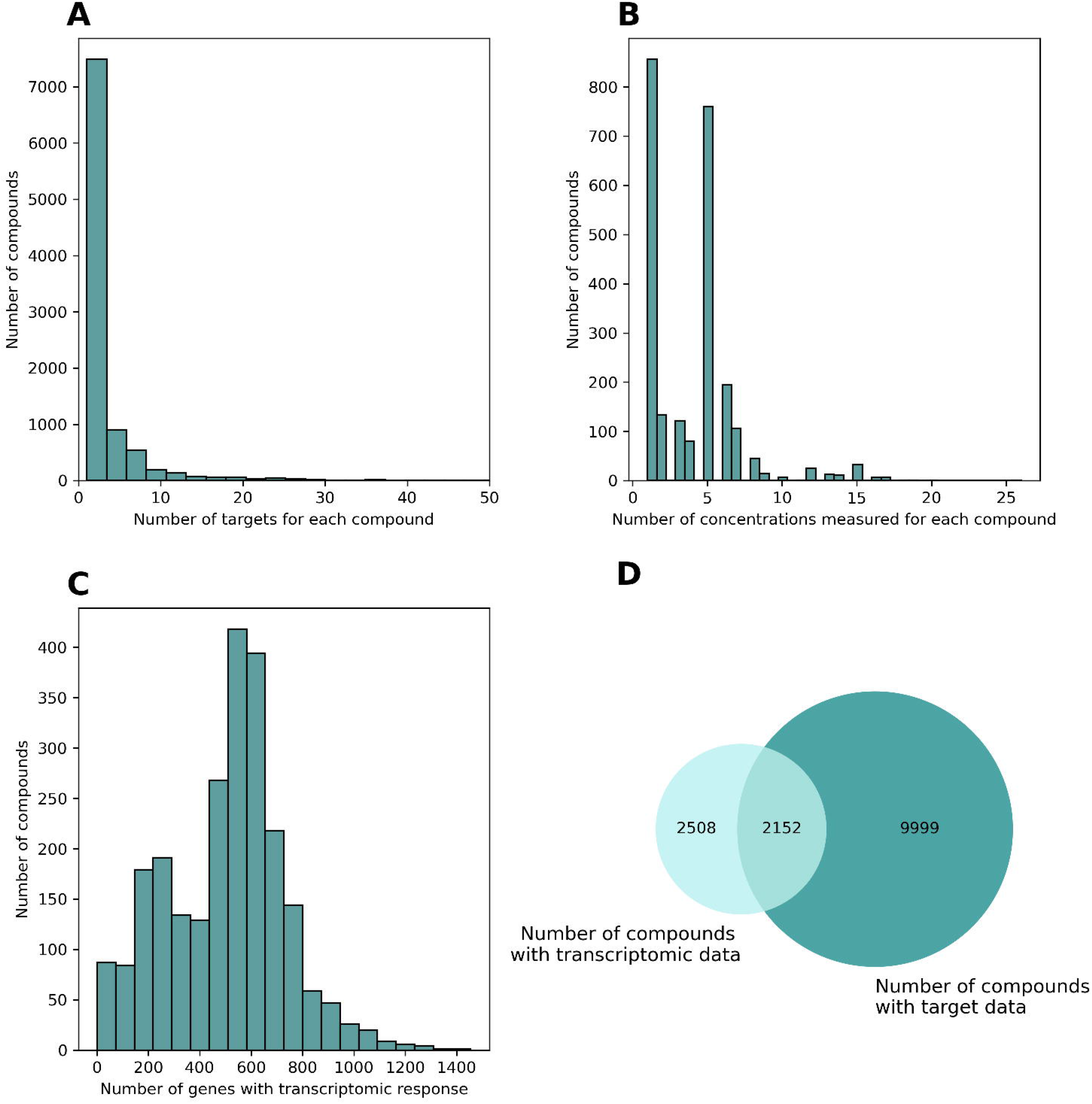

### 2.2. Representing transcriptomic and target information

After obtaining the subset of chemicals from ChemPert containing both transcriptomic and target data (i.e., chemical-target information), we represented their transcriptomic signature and their known targets as two vectors of equal length with the ultimate goal of evaluating the correlation between both. In the following subsections, we outline the two different vector representations proposed in this work.

#### 2.2.1. Original ChemPert data

Since the ChemPert data is already preprocessed and normalized, the most straightforward representation consists of directly leveraging the original vectors available in ChemPert. Thus, for each of the 2,152 compounds, we represent both its transcriptomic and target information as a vector of length X, where X corresponds to the number of genes measured (4,938) **(Figure 1)**. In both vectors, there are three possible values for each gene provided by ChemPert’s data:

- **+1**. Representing an up-regulation of the protein transcript in the case of the transcriptomic vector or activation of the protein after the chemical binds to it in the case of the chemical-target vector.
- **-1**. Representing a down-regulation of the protein transcript in the case of the transcriptomic vector or inhibition of the protein after the chemical binds to it in the case of the chemical-target vector.
- Corresponding to no change in the protein transcript or no binding

Additionally, since for a minority of the chemicals there are multiple transcriptomic experiments using different doses or concentrations **(Figure 2B)**, we represented the transcriptomic signature vector for these chemicals as the union of all differentially expressed genes. In other words, if a chemical increases the abundance of gene transcript X with a particular concentration and the same chemical decreases the abundance of gene transcript Z with a different concentration, the vector for chemical A will contain +1 for X and -1 for Z while the rest of the gene transcripts will have 0 as their value. For a small number of cases genes were both upregulated in some transcriptomic experiments and downregulated in others for a given chemical. In these cases, we first counted the number of experiments where it was upregulated and the number of experiments where it was downregulated. If it was upregulated more than two times the number of times that it was downregulated, we set the value to +1. In a similar way, if the gene was downregulated more than two times the number of times that it was upregulated, we set the value to -1. Otherwise, the vector was set to 0. While using the union allowed us to incorporate all known DEGs for a particular compound, we have additionally analyzed transcriptomic data for individual dose concentrations (e.g., dose concentration inducing the highest number of DEGs), without seeing any significant differences compared to the previously described approach **(see subsection 3.1.1)**.

#### 2.2.2. Enriching ChemPert target data with pathway information

Since compounds in the ChemPert database have only a small number of target proteins **(Figure 2A)**, the target vectors are considerably more sparse than the transcriptomic ones, meaning that it is unlikely that the two vectors will have a high degree of similarity. In order to make the target vectors less sparse, we enriched them using pathway information from KEGG, a database that contains both topological information and gene sets for over 300 pathways (Kanesha *et al*., 2021), Reactome (Gillespie *et al*. 2022), another database containing over 2,000 pathways, and WikiPathways (Martens *et al*., 2021). We denote these enriched vectors as pathway vectors through the paper. Transcriptional changes can typically be seen downstream of the target, so it is reasonable to use information about the neighbors of the target protein in the target vectors (Isik *et al*., 2015).

The process for generating the pathway vectors went as follows. For a chemical A, we obtained a target gene, which we will call gene B, by finding a gene that corresponded to a non-zero value in the target vector for chemical A. We then found all the gene sets (pathways) that contained gene B. If any of the other 4,938 genes were in any of those pathways, we changed the corresponding value of those genes in the resulting pathway vector for chemical A to match the value of gene B in the original target vector **(Figure 1B)**. We repeated this process until we had done this for every target gene of every chemical. Notably, we filtered out pathways with more than 300 genes (e.g., metabolic pathway) and pathways smaller than 15 genes, following pathway enrichment guidelines (Mubeen *et al*., 2022).

#### 2.2.3. Enriching ChemPert target data with topological information

In addition to the three pathway databases (gene sets) used above, we modified the target vectors to account for the topological information provided by KEGG. To do so, we first used the KEGG database as a directed protein-protein interaction network to be able to infer if a protein is activated or inhibited by its neighbor using the polarity of each interaction. Leveraging this network, we were able to generate protein-protein interaction vectors by modifying the values of the neighbors of each original target as follows:

- **+1** if the value of the target protein in the target vector was +1 and the target protein activated this protein or if the value of the target protein in the target vector was -1 and the target protein inhibited this protein.
- **-1** if the value of the target protein in the target vector was +1 and the target protein inhibited this protein or if the value of the target protein in the target vector is -1 and the target protein activated this protein.

If we apply these changes, we generate the values of the immediate neighbors of the target proteins in the protein-protein interaction vector. However, it is possible that these neighbor proteins also activate or inhibit other proteins farther along in the network. Thus, we also enriched the values of the neighbors in the protein-protein interaction vector by treating all of the neighbors as target proteins and repeating the above process. We repeated this process 2-5 times to test whether it would improve the correlation between the new protein-protein interaction vectors and transcriptomic vectors.

### 2.3. Correlation and Similarity Metrics

In order to calculate the agreement between a pair of vectors (x and y) for a certain compound, we used the Pearson correlation coefficient by obtaining the mean values of each vector and applying the following equation:

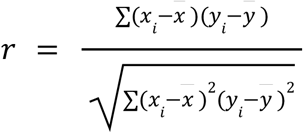

**Equation 1**. The Pearson correlation coefficient will be between -1 and 1. It is a measure of the linear correlation between the target vector and transcriptomic vector.

We also used Jaccard similarity in order to determine the similarity between pairs of vectors. We did this by applying the following equation:

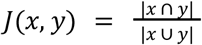

**Equation 2**. The Jaccard similarity scores are between 0 and 1 and measure the similarity between two vectors by finding the number of elements that they share.

We chose to include both metrics in our analysis in order to be more transparent about the results. Since the data in our vectors is discrete, Jaccard similarity allowed us to measure how many of the values matched up across the two vectors. Complementary, Pearson correlation allowed us to see if the two vectors were positively or negatively correlated.

### 2.4. Correlation Analyses

#### 2.4.1. Correlating the transcriptomic and target vectors for a given compound

We applied the aforementioned correlation metrics (subsection 2.3) in order to calculate the agreement between a target vector, x, and a transcriptomic vector, y, for a certain compound **(Figure 1A)**. After calculating the correlation coefficients for each compound, we used the permutation test to determine whether the results were statistically significant. We permuted the pairs of target vectors and transcriptomic vectors and applied the same correlation metrics to these random pairs. We repeated this process 100,000 times and obtained the mean correlation score for each iteration. We then found the *p*-value by using the distribution of correlation means from the permutation tests We used a significance level of 0.05 to determine whether our results were statistically significant after applying Bonferroni correction.

#### 2.4.2. Investigating pairs of compounds based on their transcriptomic/target vector similarity

Apart from assessing whether the transcriptomic and target vectors of a given compound have any correlation, we sought to evaluate whether there was any correlation between the target vector similarity scores and the transcriptomic vector similarity scores. In order to do this, we first calculated the Jaccard similarity for every pair of transcriptomic vectors and every pair of target vectors. To discard the inherent variability across cell lines, we filtered out compound pairs that were not tested in the same cell line. We then created a vector, which we call X, of all of the target vector correlation scores and another vector, which we call Y, of all the transcriptomic vector correlation scores. We ensured that for each compound pair the target similarity score and transcriptomic similarity score corresponding to that pair were in the same position in their respective vectors. In order to determine whether target vector similarity was correlated with transcriptomic vector similarity, we calculated the Pearson correlation coefficient for X and Y **(Figure 1C (3))**. To determine whether the Pearson correlation coefficient was significant, we used the permutation test by permuting the transcriptomic similarity vector, Y, 100,000 times.

In order to better understand whether pairs of compounds with similar transcriptomic profiles also shared targets, we first calculated the Jaccard similarity (equation 2) for every pair of transcriptomic vectors **(Figure 1C (2))** and filtered out any pairs with a correlation coefficient less than 0.6. The rationale behind choosing 0.6 as a cutoff was we wanted to include a large sample of compound pairs in our study and there were a limited number of compound pairs (32) with a transcriptomic signature correlation score above 0.7. After selecting these pairs, we calculated the Jaccard similarity of the target vectors for each pair and used those correlation coefficients to determine whether compounds with similar transcriptomic profiles shared targets.

Following, we decided to evaluate whether compounds with the same targets had similar transcriptomic profiles measured in the same cell line. To this end, we calculated the Jaccard similarity scores for all the target vector pairs **(Figure 1C (2))** and filtered out any compound pairs whose correlation score was less than one (i.e., all the target(s) of a compound perfectly match the targets of the other one). We employed such a strict cutoff because we only wanted to examine pairs that shared all of their targets and affected those targets in the same way (up-regulation or down-regulation). For the remaining pairs, we calculated the Jaccard similarity scores (equation 2) for their transcriptomic vectors and utilized those correlation coefficients to determine whether the compound pairs with shared targets had similar transcriptomic profiles.

#### 2.4.3. Permutation analysis at pathway level

We also used a similar method to determine the statistical significance of the correlation scores between the transcriptomic vectors and the target vectors that were modified using pathway information. In order to do this, we first removed any pathways from the original pathways that contained more than 300 genes or less than 15 genes. We found the lengths of the remaining pathways in the dataset and filled the pathways up with random genes from the pathways. We did this process 1,000 different times. We then calculated the mean Pearson correlation score and Jaccard similarity score for each set of random pathways and compared these scores to the mean correlation scores for the original pathways.

#### 2.4.4. Permutation analysis at network level

In order to further determine the statistical significance of the correlation scores between the target vectors that were modified with topological information from KEGG and the transcriptomic vectors, we generated permuted networks by using the XSWAP algorithm (Hanhijärvi *et al*., 2009). We then created a new set of target vectors for each random network using the method from subsection 2.2.3. After creating these vectors, we calculated the mean Jaccard similarity score and the mean absolute value of the Pearson correlation score between the target vectors and transcriptomic vectors for each network and compared these scores to the scores obtained with the original network. In this case, we used the mean absolute value of the Pearson correlation scores because we were more interested in knowing whether the vectors were correlated than knowing if they were positively or negatively correlated.

### 2.5. Implementations details and code availability

We preprocessed the datasets and generated the vectors using the Pandas Python package (McKinney, 2010). In the enrichment analysis, we modified these vectors by leveraging the KEGG pathways available at PathMe (Domingo-Fernández *et al*., 2019) (released date 01-03-2021). To calculate the correlations, we employed the NumPy (Harris *et al*., 2020) and SciPy (Virtanen *et al*., 2020) Python packages. Additionally, we plotted the visualizations using Seaborn (Waskom, 2021) and Matplotlib (Hunter, 2007). Lastly, both data and source code are available at https://github.com/enveda/transcriptomic-target-correlation.

## 3. Results

### 3.1. Targets are generally not differentially expressed in drug perturbation transcriptomic experiments

We first investigated the correlation between the target vectors and transcriptomic vectors retrieved from ChemPert without any modification. The resulting correlation scores were very low with the majority of the Pearson correlation scores between -0.02 and 0.02 **(Figure 3A)**. This was not a surprising result for two reasons. Firstly, the target vectors are very sparse **(Figure 1A)** compared to the transcriptomic vectors **(Supplementary Figure 2)**. Furthermore, most of the known targets are not differentially expressed genes **(Supplementary Figure 1)**, in line with previous work (Isik *et al*., 2015). In an effort to determine whether the correlation scores were statistically significant, we used permutation tests on both the target and transcriptomic vectors and compared the results against the null distribution. This analysis confirmed that the permuted vectors yielded comparable Pearson correlations to the original ones (*q*-value = 1.54) **(Figure 3B)**. Similarly, the Jaccard similarity scores obtained on the permuted vectors were not significantly different (*q*-value = 2.412).

**Figure 3.**
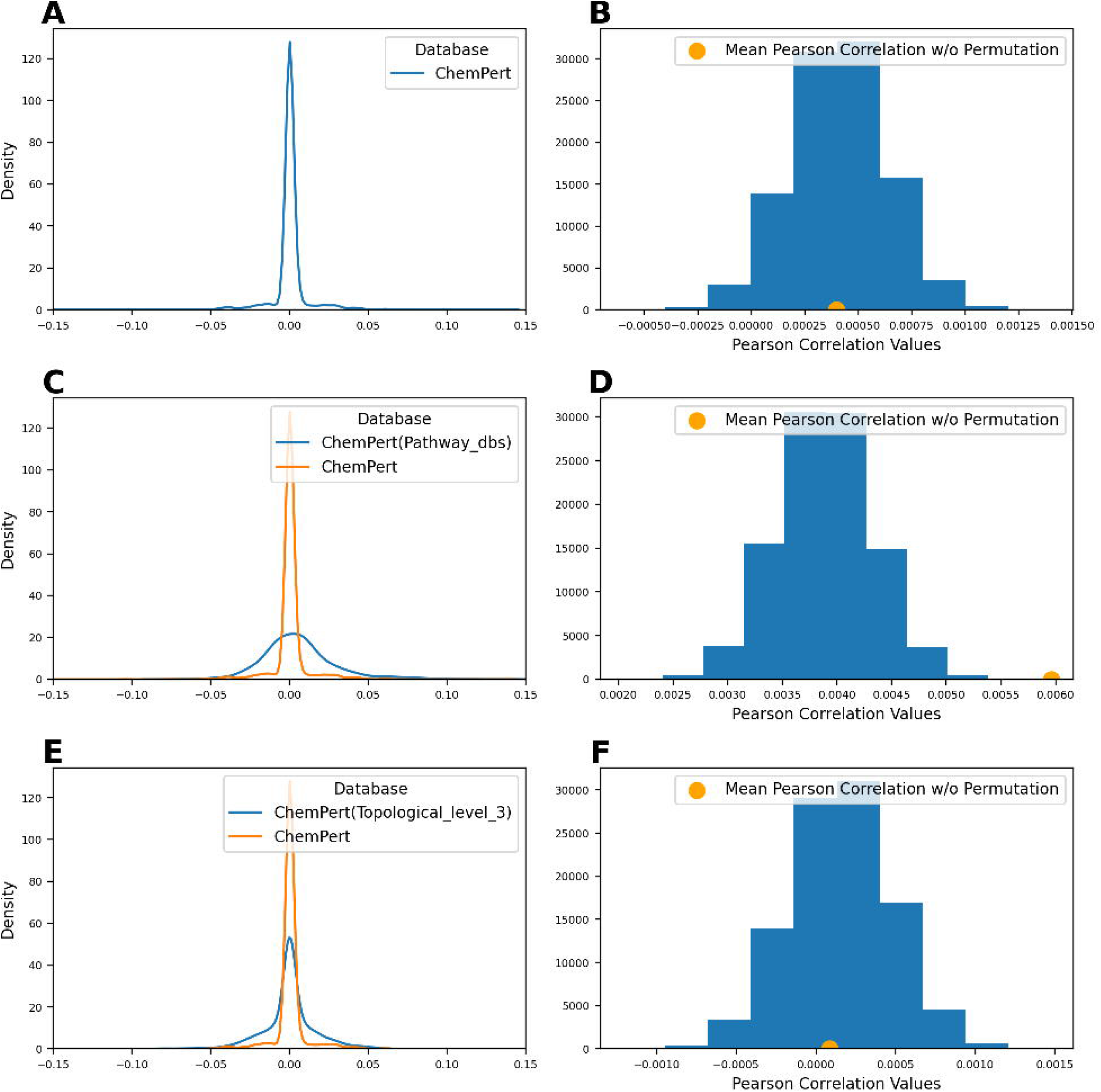

#### 3.1.1. Using transcriptomic responses with highest number of DEGs yielded similar results

ChemPert reported several transcriptomic responses for a minority of the compounds in our dataset. In addition to representing the transcriptomic vectors as a union of the differentially expressed genes (see subsection 2.2.1), we also tried using the transcriptomic response with the highest number of DEGs for each compound to create the transcriptional vectors. When we calculated the correlation between the target vectors and these transcriptomic vectors, the majority of the Pearson correlation scores were between -0.02 and 0.02 (similar to the results in section 3.1). The Jaccard similarity scores for these vectors were also similar. Since the results for both sets of transcriptomic vectors were not significantly different, we decided to use the transcriptomic vectors that combined the different responses for the rest of the analysis.

### 3.2. Drug perturbation transcriptomic signatures tend to correlate with the downstream pathways targeted

After observing the low correlation coefficients for the ChemPert target vectors and transcriptional vectors, we hypothesized that enriching the target vectors with downstream pathway information might increase our correlation scores due to two main reasons: i) transcriptomics signatures capture downstream changes at the pathway level (Garrido-Rodríguez *et al*, 2022, Isik *et al*., 2015), and ii) the pathway vectors are less sparse compared with the target vectors **(Supplementary Figure 3B)**. Below, we discuss the results after conducting two disparate enrichment approaches.

#### 3.2.1. Pathway level analysis

The first and simplest enrichment approach consisted of using pathway vectors corresponding to proteins that participate in the same pathway as the original target(s) (**Figure 1B**). Conducting this enrichment increased the average Pearson correlation scores between target and transcriptomic vectors from 0.00039 to 0.00596 compared to the original ChemPert correlation scores **(Figure 3C)**. While these initial results appeared to be positive, we wanted to make sure that these results were statistically significant and not a result of the pathway vectors being less sparse. Thus, we conducted 100,000 permutations experiments on the transcriptomic vectors and pathway vectors and compared their underlying distribution of Pearson correlation scores to the observed one (*q*-value of 0.00003 **(Figure 3D)**) Similarly by comparing the Jaccard similarity scores for these permutation tests to the null distribution, we got a *q*-value of 0.09846.

Additionally, we wanted to ensure that the pathways from KEGG, Reactome, and WikiPathways were not only increasing the correlation scores because the enriched target vectors were less sparse than the original target vectors. Thus, we generated 1,000 sets of random pathways maintaining their original size and gene occurrence and enriched the target vectors with each set of random pathways. We compared the mean similarity scores and correlation scores for these random sets of pathways to the scores for the original set of pathways. The mean Pearson correlation score and mean Jaccard similarity score for the original pathways were significant (*q*-value = 0.003). Thus, we determined that the higher correlation scores resulting from enriching the target vectors with pathway information were not just a result of making the target vectors less sparse.

#### 3.2.2. Network level analysis

The second enrichment approach leverages topological information from KEGG to generate protein-protein interaction vectors. Using the protein-protein interaction vectors increased the Jaccard similarity scores. Furthermore, the similarity scores continued to increase when we repeatedly applied the database up to three levels downstream of the target **(Figure 3E)**. When we used the permutation test on this data, the results were not statistically significant up to level two. This is not an unexpected result since the vectors were only slightly enriched at these first two levels. However, after the enrichment with topological information going three levels downstream of the target, our results were statistically significant using Jaccard similarity (*q*-value of 0.00015) **(Figure 3F**). We also calculated the Pearson correlation score between the protein-protein interaction vectors and the transcriptomic vectors. However, the Pearson correlation score did not increase and our results were not statistically significant when we used Pearson correlation (*q*-value of 1.772). Thus, indicating the importance of conducting the correlation/similarity analyses with non-sparse vectors.

In order to determine whether the topological information from KEGG was significant, we generated 100 random networks of the same structure. We then enriched the target vectors with the information from each of the random networks (subsection 2.4.3). We then computed the correlation scores and similarity scores between these sets of enriched target vectors and the transcriptomic vectors. When we compared these scores to the scores of the original network, the original network had a *q*-value of 0.15 when we used Jaccard similarity and had a *q*-value of 0.03 when the used the mean absolute value of the Pearson correlation scores. Thus the topological information from KEGG added significant information to the target vectors.

In summary, our results indicate that enriching target vectors with pathway information increases their correlation with transcriptomic signatures. This is not surprising given that previous work observed that differentially expressed genes tend to be closer to the target than randomly expected (Isik *et al*., 2015).

#### 3.2.3. Assessing the impact of pathway size

Since large pathways are more likely to have a non-zero value in the vector, we evaluated whether removing large pathways had an influence on the observed correlations. To do so, we generated different subsets of the pathway dataset where the top X largest pathways were removed (i.e., top 50, top 100, top 250) as well as subsets of pathways with varied numbers of genes. We used the subsets to enrich the target vectors and calculated the correlation scores between these target vectors and the transcriptomic vectors. The correlation scores were similar for all subsets of the pathways **(Supplementary Figure 4)**. We also applied permutation tests to these vectors and found that all sets had a *q*-value of 0.003 when we used Pearson correlation. When Jaccard correlation was used, the *q*-values ranged from 0.057 to 0.27 with the target vectors using the set of pathways with 100-300 genes having the highest *q*-value.

### 3.3. Target and transcriptomic vector similarity are slightly correlated

Our next question was whether compounds with similar targets had similar transcriptomic responses or vice versa. To discard the inherent variability across cell lines, we filtered out compound pairs that were not tested in the same cell line. We then calculated the Jaccard similarity scores between each target vector pair and each transcriptomic vector pair to identify pairs of compounds targeting the same proteins. In order to determine whether the target similarity scores were correlated to the transcriptomic similarity scores for each compound pair, we created x,y coordinate pairs of the target and transcriptomic similarity scores and computed the Pearson correlation coefficient for all of these pairs **(see Methods; Figure 1C (3))**. When we did this with the vectors from the ChemPert database, we obtained a Pearson correlation coefficient of 0.012. We used a permutation test with 1,000 permutations and determined that this correlation coefficient was significant with a significance level of 0.05 (*q*-value of 0.003). We also repeated this process after we had enriched the target vectors with pathway information and we computed a Pearson correlation coefficient of 0.052 which was also statistically significant (*q*-value of 0.003). To summarize, these results reveal that there is a slight correlation between the target similarity for a compound pair and the transcriptomic similarity for that same compound pair.

#### 3.3.1. Compounds with similar transcriptomic profiles are more likely to share targets

We decided to further investigate whether compounds that induced similar transcriptomic profiles target the same proteins by filtering out those pairs with a transcriptomic correlation score less than 0.6 **(Figure 4**) **(Supplementary File 1)**. Among these over 240 pairs, the average target correlation score was 0.044 compared to the correlation scores for all possible compound pairs which was 0.005. Thus, the average target correlation score did increase when we only looked at pairs with high transcriptomic similarity. Interestingly, the percentage of compound pairs that shared at least one target also increased from 2.88% to 18.85%. Additionally, when we increased the cutoff for transcriptomic signature similarity from 0.6 to 0.7, the percentage of pairs with at least one shared target increased to 34.3% (11/32). Furthermore, all of the compound pairs (6) with transcriptomic signature similarity scores over 0.8 share at least one target. For instance, the pair with the highest correlation is Alvocidib and Cgp-60474, both of which are Cyclin Dependent Kinase (CDK) inhibitors. Alvocidib is a flavonoid alkaloid CDK9 kinase inhibitor extracted from two plants and CGP60474 a potent CDK inhibitor. Taken together, our findings suggest that pairs of chemicals which have a high correspondence on the transcriptomic vector tend to share at least one target.

**Figure 4.**
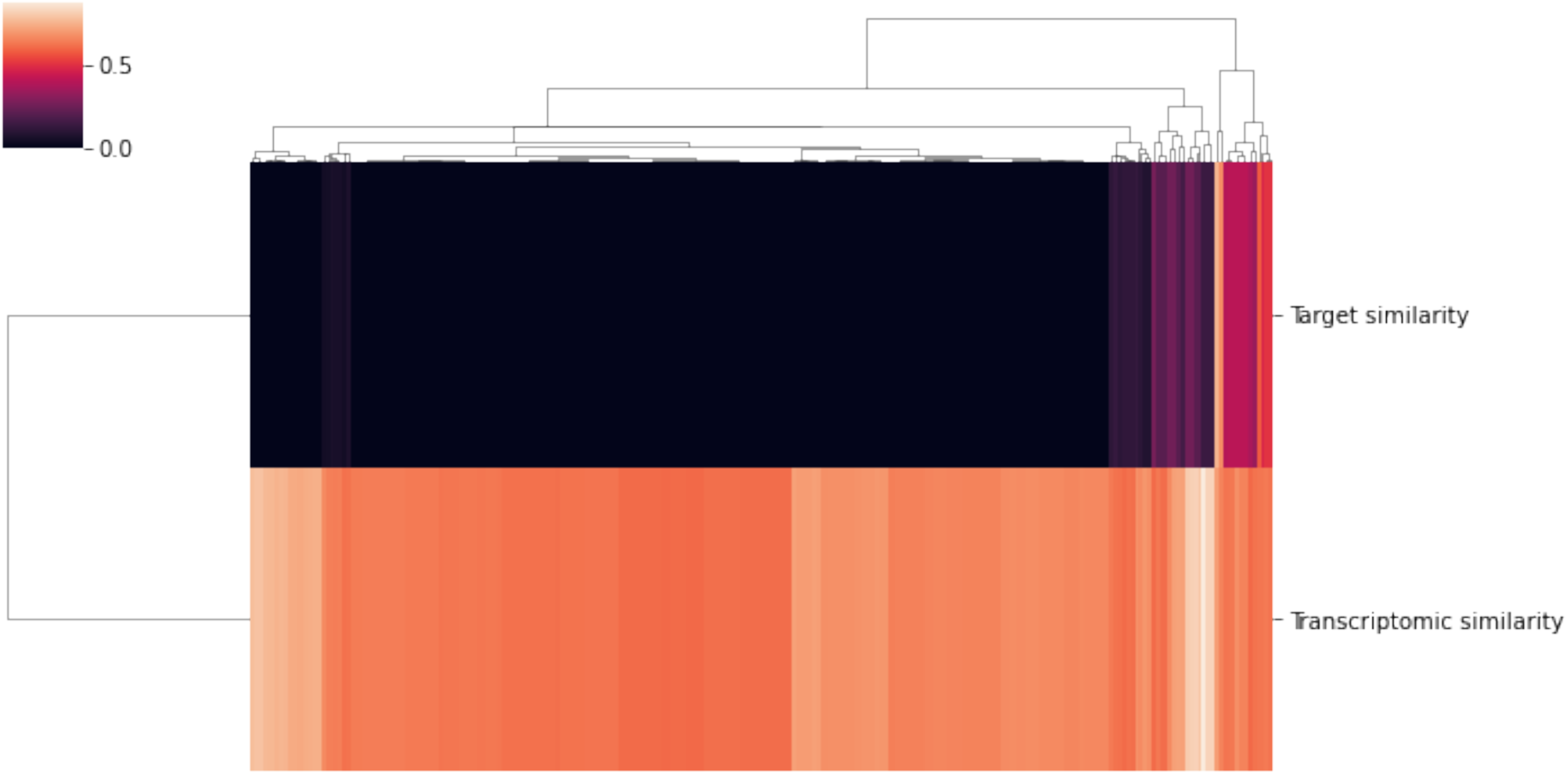

#### 3.3.2. Compounds targeting the same targets typically induce disparate transcriptomic responses

Similarly, we also decided to evaluate whether compounds with the same targets induced similar transcriptomic responses. In order to find the compounds with the same targets, we calculated the Jaccard similarity for all possible pairs of target vectors and filtered out any compound pairs whose correlation score was less than one. After identifying those compound pairs that share targets (2,435) **(Supplementary File 1)**, we calculated their Jaccard similarity using their transcriptomic vectors. Interestingly, the average transcriptomic correlation score was 0.139 which we compared to the average correlation score of 0.130 for all possible compound pairs. Furthermore, we found that all the transcriptomic correlation scores were lower than 0.6, the threshold previously used in subsection 3.3. This suggests that even when compounds share all of their target(s), they do not necessarily induce a similar transcriptomic response.

Despite the low correlation observed between transcriptomic signatures of compounds sharing targets, we investigated the eight pairs of compounds that had a correlation equal to or higher than 0.5. Among these, we found four pairs of natural products with well-known uses for treating arrhythmias among other indications (i.e., digitoxigenin, sarmentogenin, cymarin, proscillaridin). These compounds are structurally related and are cardenolides, a family of heart poison compounds from plants. The highest correlated pair is digitoxigenin and sarmentogenin **(Figure 5)**. Both are used due to their ability to inhibit ATP1A1 (El-Seedi *et al*., 2019). Additionally, we found a pair of DNA topoisomerase I inhibitors (i.e., Genz-644282 and SN-38), a pair of JNK inhibitors (i.e., ZG-10 and JNK-9L), and a pair of dual PYK2/FAK inhibitor (i.e., PF-431396 and PF-562271).

**Figure 5.**
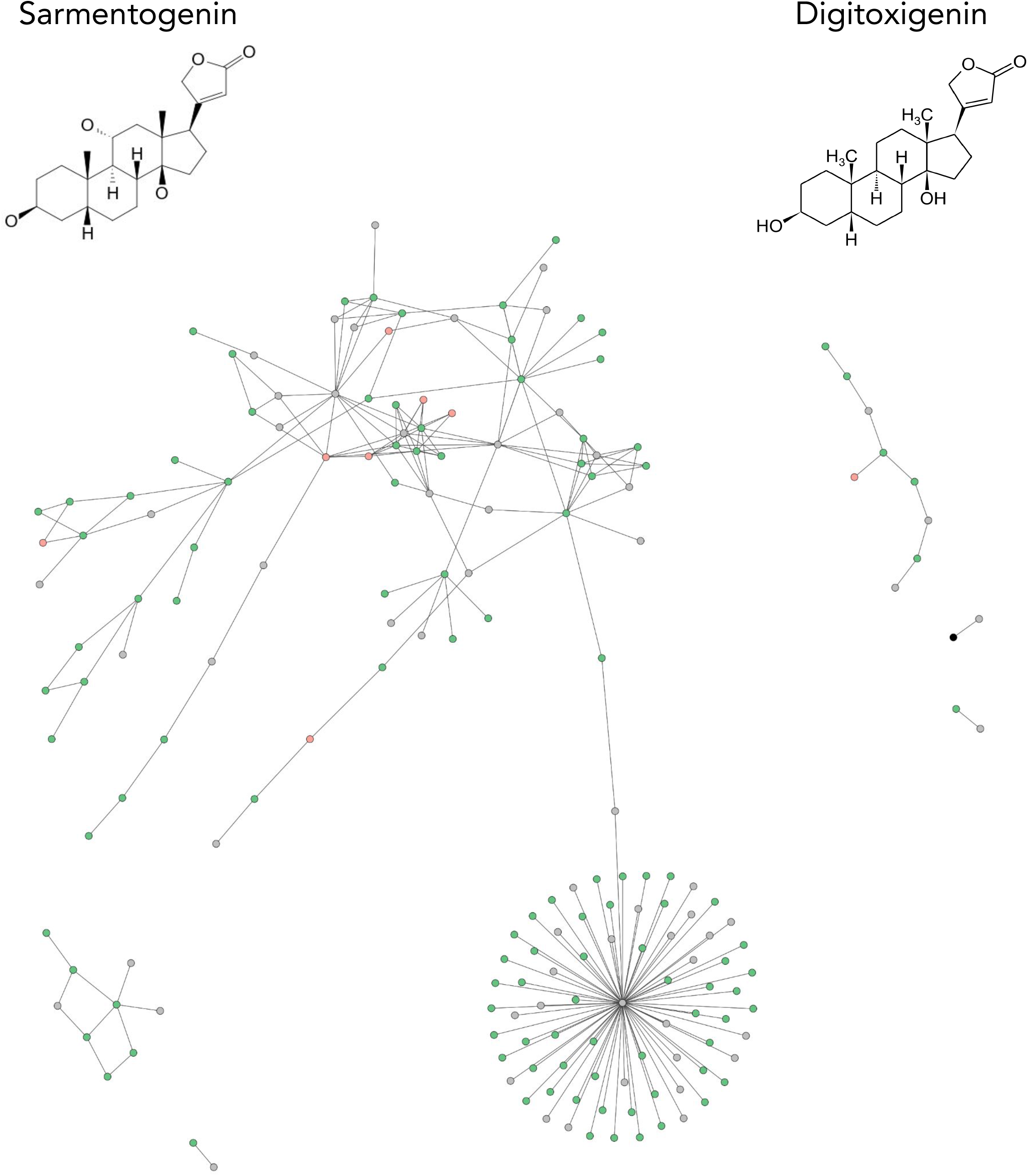

## 4. Discussion

In this work, we systematically evaluated correlations between target and transcriptomic data for 2,152 compounds. In line with prior work, we found that target and transcriptomic signatures do not correlate, as targets are rarely differentially expressed in transcriptomic experiments. However, once target information is enriched with pathway information, the correlation between both increases. Additionally, we investigated whether compounds that target the same proteins induce a similar transcriptomic response. We found that pairs of compounds sharing the same target show a slightly increased transcriptomic correlation than average. Inversely, we found that compounds with similar transcriptomic profiles are more likely to target at least one shared protein and often treat the same indication. Lastly, we focused on two natural products (i.e., sarmentogenin and digitoxigenin, which are two members of the cardenolide family) that share all their target proteins, and also have the highest transcriptional similarity in order to demonstrate how exploring their high correlations can elucidate their MoA.

Nonetheless, the presented work is not without its limitations. First, our analysis could have been complemented by leveraging the raw transcriptomic profiles, as we have only employed the transcriptomic profiles already processed by ChemPert. This would have allowed us to lower the significance threshold and assess whether the threshold plays a role in the observed correlations between transcriptomic and target vectors. However, it would require significant effort as thousands of experiments would have to be processed. Second, since some compounds have transcriptomic profiles for multiple concentrations, one can take several approaches to correlate the profiles (e.g., use the one with the largest number of DEGs, the one with the highest concentration, etc.). While our main results were generated by taking the union of all differentially expressed genes in these profiles, we also tested alternative approaches using the profile with the largest number of DEGs and observed similar results (subsection 3.1.1). Third, it is important to note that the transcriptomic profiles were obtained from different cell lines. While it is well-known that gene expression patterns vary among cell lines (Figueiredo *et al*., 2022), Pabon *et al*. (2018) found that the cell line exhibiting the lowest correlation with respect to the control yielded the best results for target prediction. Fourthly, while we could only employ a subset of ChemPert (i.e., over 2,000 compounds), this is still a larger number of profiles than previous studies that investigated transcriptomic correlations (Pabon *et al*., 2015; Isik *et al*., 2016). Lastly, the data we leveraged from ChemPert was conducted on the L1000 platform which infers the expression of 11,350 genes from 978 measured genes. This, combined with the fact that only 4,938 of these genes were targets for any of the 2,152 compounds, restricted the protein space of our exploration since not all genes/proteins were represented in the vectors. However, this set of approximately 5,000 genes that was used to conduct this study highly overlap with the genes included in pathway databases; thus, indicating that they correspond to the ones for which we have more functional information about.

In the future, we ambition multiple possible extensions of our work. Firstly, a prospective study could validate our findings by leveraging an additional database. Secondly, additional *omics* modalities beyond transcriptomics such as proteomics and metabolomics could be incorporated in our analysis to explore whether higher correlations are observed. Thirdly, we could assess if the correlations between the transcriptomic and target information tend to be higher when the transcriptomic experiment has been measured in a cell line characteristic to the particular tissue where a drug acts (e.g., neuron or glial cells for drugs treating neurological disorders). If this would be the case, these higher correlations could be used as a proxy to identify candidate repurposing drugs (Wagner *et al*., 2015; Namba *et al*., 2022). Alternatively, as demonstrated in our case study, correlating both transcriptomic and target information can be used to better understand the MoA of a drug. Additionally, one can use the transcriptomic or target vectors as compound features and conduct a classification task using machine learning models to, for instance, predict the type of compound, their targets, etc.. Finally, another possible application to the problem of repurposing compounds would be to infer novel activities of combinations of compounds by concatenating their transcriptomic profiles.

## Authors’ Contributions

CEH implemented and performed the analyses. CEH and DDF prepared and generated the datasets and networks. CEH, DE and DDF interpreted the results. DDF conceived the study. DE and DDF designed the study. DE, DH and DDF supervised the study. CEH, DE and DDF wrote the paper.

All authors have read and approved the final manuscript.

## Acknowledgements

We would like to thank the entire ChemPert team for releasing the data that was used as the foundation of this work.

