## Supplementary for "On the correspondence between the transcriptomic response of a compound and its effects on its targets"

### Supplementary Figures

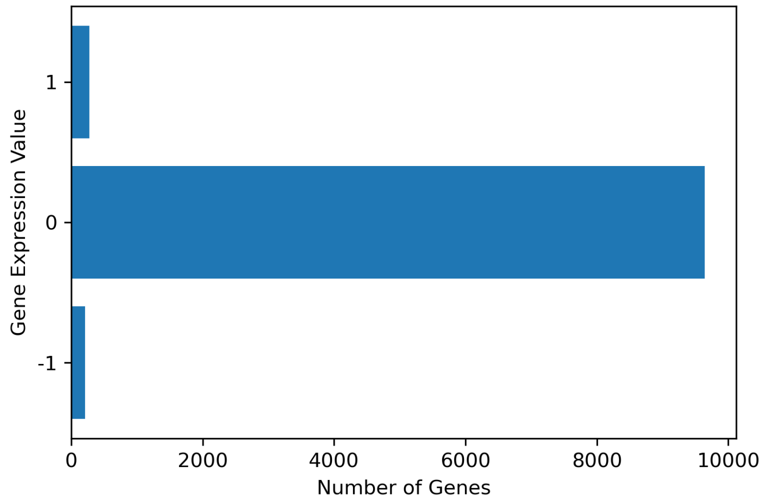


**Supplementary Figure 1.** Gene expression values for known targets in the ChemPert database.

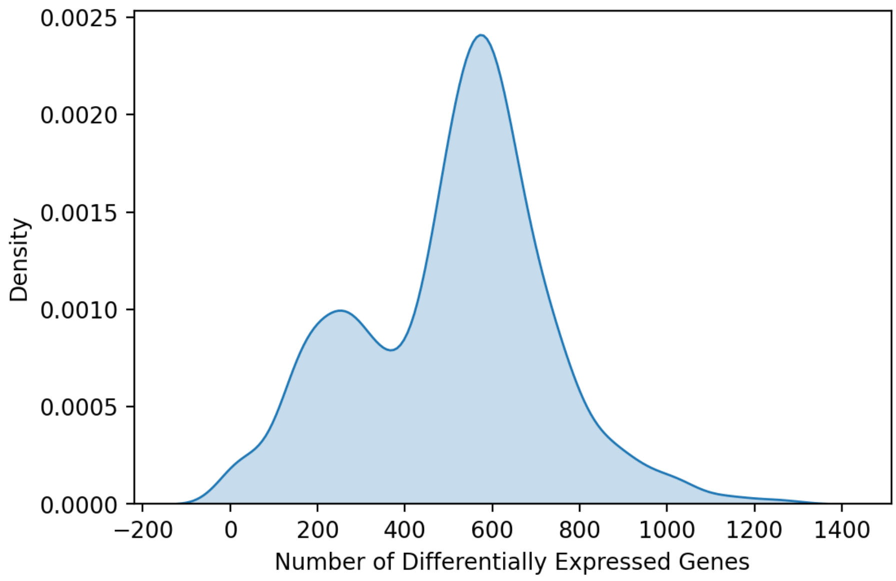


**Supplementary Figure 2.** Distribution of differentially expressed genes for each chemical

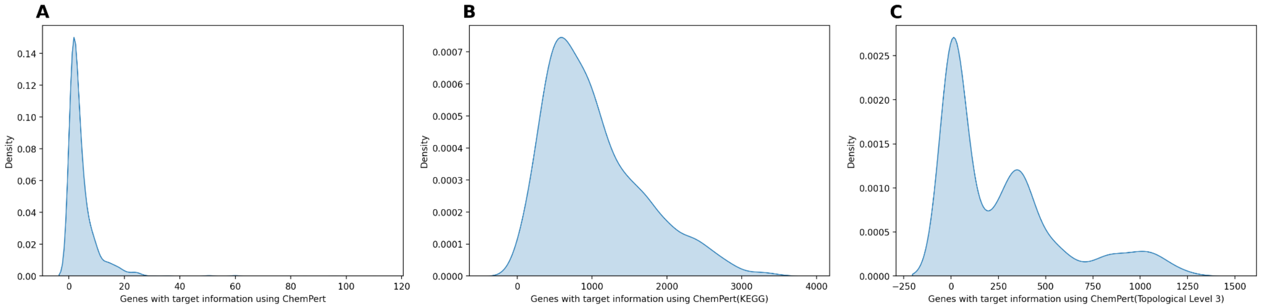


**Supplementary Figure 3. a)** Distribution of targets for each chemical using the original data from the ChemPert database. b) Distribution of targets for each chemical after enriching them with the gene sets from the KEGG database. c) Distribution of targets for each chemical after the enrichment using topological information up to three levels downstream of the target(s).

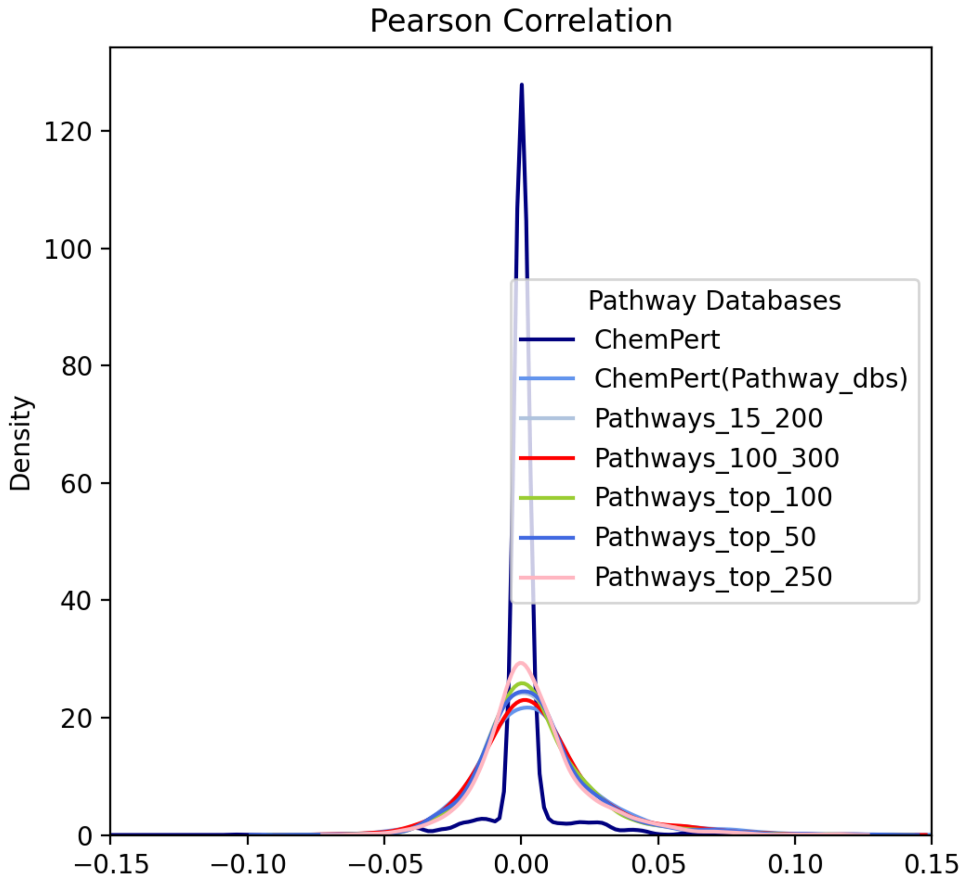


**Supplementary Figure 4.** Pearson correlation scores for pathway and transcriptomic vectors from ChemPert for each of the 2,512 compounds using different subsets of pathways. We first tried keeping pathways from the dataset with 100-300 genes (Pathways_100_300). When we did this, we were only left with 300 pathways (1660 pathways are kept when we use pathways with 15-300 genes). Additionally, we tried keeping pathways with 15-200 genes. This resulted in a set of 1600 pathways. We also tried removing the pathways with the top 50,100, and 250 number of genes (Pathways_top_X where X is the number of pathways we removed). We used each of these sets of pathways to adjust the target vectors (as described in section 2.2.2) and computed the Pearson correlation and Jaccard similarity between these new target vectors and the transcriptomic response vectors.

### Supplementary Text

The ChemPert database includes 82,270 transcriptional signatures from 167 different cell types. The data we used was generated from 2,508 unique perturbagens. They collected data about the targets of different perturbagens from Drugbank, STITCH, and Drug repurposing Hub. The perturbagens that were used included both chemical and biological perturbagens. Information about the gene expression of different cell types and the transcriptional responses was collected from Gene Expression Omnibus (GEO), ArrayExpress, and LINC L1000. The dataset of transcriptional responses was manually curated and included responses from non-cancerous cells in humans, mice, and rats. For more information about how the data was collected and processed see the original work (Zheng *et al.*, 2022).
